# Hemispheric multi-dimension features extraction analysis based on decoupled representation learning

**DOI:** 10.1101/2024.03.13.584299

**Authors:** Yuwei Su, Sifeng Wang, Xiaoyu Zhang, Min Lan, Suyu Zhong

**Author notes:** Address correspondence to Dr Suyu Zhong, School of Artificial Intelligence, Beijing University of Posts and Telecommunications, #10 Xitucheng Road, Haidian District, Beijing 100876, China.

## Abstract

The predominant approach in investigating brain structural asymmetry relies on predefined regions of interest, assessing variations between homologous brain regions through a single indicator, which is local, univariate, and relative. In response to this challenge, we employ decoupled representation learning from deep learning to extract hidden features containing hemisphere-specific information at a hemispheric systemic level. This novel approach enables a global and multivariate analysis of brain structural asymmetry. Our findings indicate a significant association between left-hemisphere-specific hidden features and language-related behavioral metrics, as well as a correlation between right-hemisphere-specific hidden features and social-related behavioral metrics. Tensor-based Morphometry results find the impact of left-hemisphere-specific features on the left inferior frontal sulcus within Broca’s area, a crucial region for language processing. Additionally, right-hemisphere-specific features influenced the right rostral hippocampus, a region implicated in emotion regulation and spatial navigation. The findings from Neurosynth indicate that significant regions caused by left-hemisphere-specific features are correlated with language, while significant regions caused by right-hemisphere-specific features are associated with behaviors primarily governed by the right hemisphere. Furthermore, our study establishes a link between structural changes induced by hemisphere-specific features and several genes. Such findings demonstrate that the application of deep learning techniques allows for precise capture of hemisphere-specific information within individual hemispheres, offering a new perspective for future research on brain structural asymmetry.

## 1. Introduction

Since Broca’s groundbreaking identification of a specialized region in the left frontal lobe associated with language [1], the past few centuries have witnessed attention-grabbing studies on human brain asymmetry [2,3,4,5]. As of now, it is well-established that the left hemisphere plays a primary role in language abilities [6,7,8,9], while the right hemisphere may excel in tasks related to visual-spatial processing, face recognition, and social cognition [9,10,11]. This functional specialization across hemispheres, believed to reduce redundancy in brain functions, could optimize computational and energetic efficiency [12].

In addition to researches on functional asymmetry, there are also extensive studies on structural asymmetry that holds significant implications for understanding neuroplasticity, neurodevelopmental and psychiatric disorders. Steinmetz found that patients with developmental deficits of phonological processing showed decreased planum temporale asymmetry, while musicians displayed exaggerated leftward asymmetry, which revealed the correlation between changes in brain structure asymmetry and alterations in brain function [13]. Postema pointed altered asymmetries of cortical thickness in medial frontal, orbitofrontal, inferior temporal, and cingulate regions in autism spectrum disorder compared with controls [14]. Changes of brain structural asymmetry have also been found in other disorders including schizophrenia, obsessive–compulsive disorder and Alzheimer’s disease [15,16,17].

The asymmetry index (AI), calculated by dividing the difference of a specific measure, i.e., cortical thickness, between homologous brain regions in the left and right hemispheres by their sum, is a prominent metric for investigating cerebral structural asymmetry [5]. However, there exists some controversy regarding AI. For example, Williams emphasized that experts have not explicitly clarified why dividing by the sum of left and right hemisphere measures in the calculation of AI is necessary [18]. Furthermore, AI only considers the difference in a single measure between homologous brain regions, making the asymmetry of each brain region pair independent of others [19], which may result in the loss of information concerning interactions between various brain regions. Also, examining only one measure at a time would limit researchers with overly simplified assumptions, making it challenging to capture the complex relationships existing within the brain system.

In addressing the aforementioned challenges, numerous scholars have proposed new avenues of research. Liu asserted that lateralized brain systems resulted from multiple distinct factors [20]; Kong proposed that brain asymmetry should be viewed as a complex and multivariate trait [21]; Saltoun decomposed the asymmetry of brain structure into different patterns [19]. Apart from innovations in research perspectives, neuroscientists have also employed new methodologies. For instance, applying a branch of artificial intelligence, i.e., decoupled representation learning which aims to untangle the potential generating factors across multiple levels and scales within the data, Jung-Hoon Kim disentangled latent representations of resting-state functional images to examine the spatiotemporal characteristics of resting-state functional activity [22]; Aglinskas disentangled specific features in brain structural images among individuals with Autism Spectrum Disorder (ASD), differentiating them from features shared with healthy controls [23].

Inspired by these studies, we hypothesized that the human brain could be unraveled into three distinct factors: left-hemisphere-specific factors, right-hemisphere-specific factors, and factors shared by both hemispheres. We conceived and implemented a decoupled representation learning model to distill distinct left-hemisphere-specific, right-hemisphere-specific, and shared features from human anatomical hemispheres. Left-hemisphere-specific features and shared features combined to form the left hemisphere, while right-hemisphere-specific features and shared features combined to form the right hemisphere. Building on this conceptual framework, we could examine the systematic level of hemispheres within the human brain to avoid the loss of potentially significant information and investigate structural asymmetry from a multivariate perspective by considering different factors. Specifically, we leveraged structural magnetic resonance imaging (MRI) from Human Connectome Project (HCP) to train our model and extract hemisphere-specific features. Then we validated whether the extracted hemisphere-specific features contained hemisphere-specific information using partial least squares correlation (PLSC). Subsequently, we employed the tensor-based morphometry (TBM) method to identify brain regions exhibiting significant structural changes caused by hemisphere-specific features. Additionally, we conducted a meta-analysis to investigate the relationship between significant changed brain regions and cognitive behavioral metrics. Finally, a gene enrichment analysis was performed. The workflow of this article was presented in Figure 1.

**Fig. 1.**
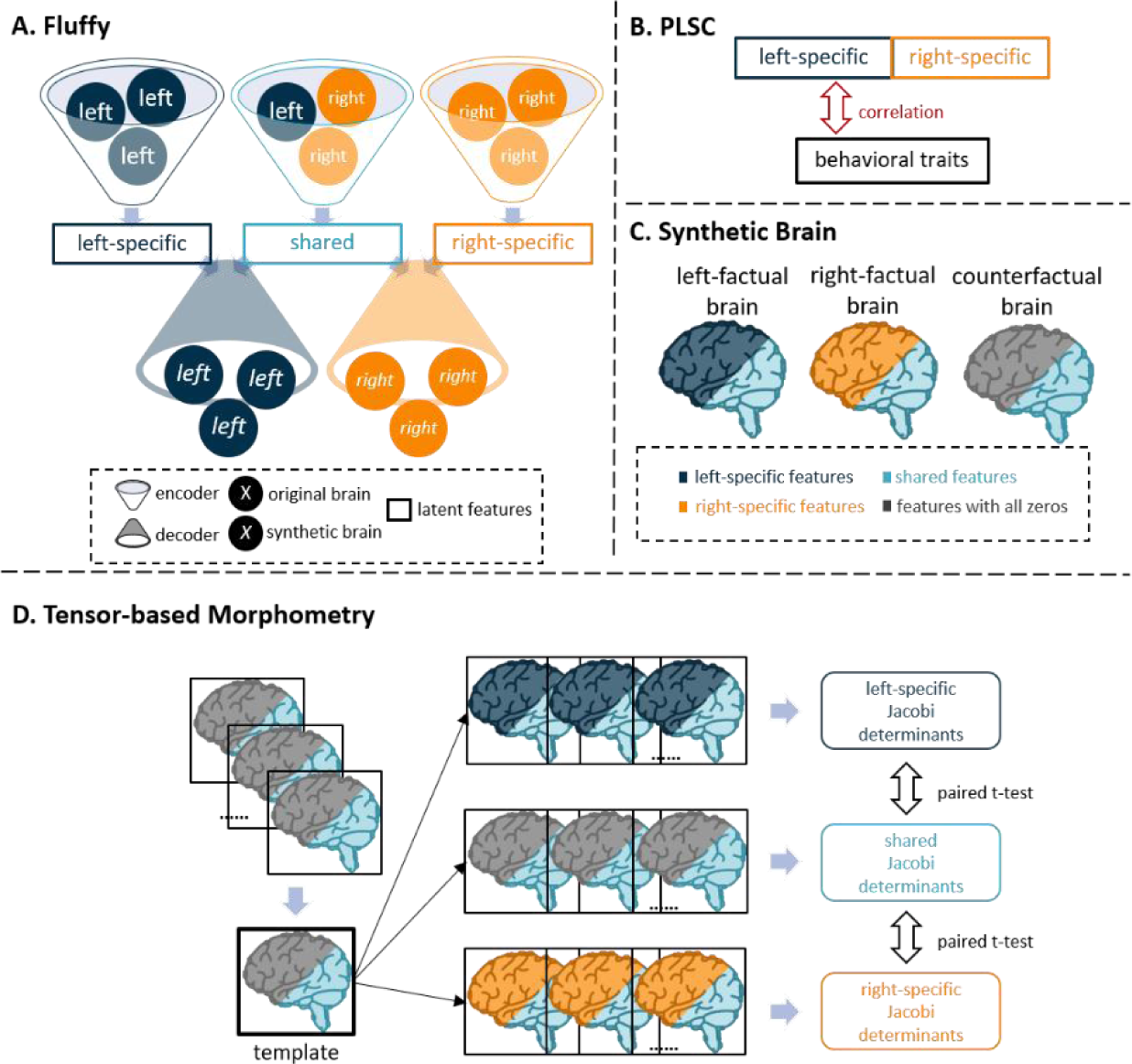
Workflow. (A) Decoupled representation learning model called Fluffy. The three encoders took separated left- and right-hemisphere brain as input, yielding three sets of latent features. The two decoders received a combined input of hemisphere-specific and shared features, generating synthetic hemisphere data. (B) PLSC was performed to identify interpretable relationships between hemisphere-specific features and widely recognized behavioral traits favoring one hemisphere (language, social). (C) Two factual and a counterfactual brains were produced for each subject. ‘Features with all zeros’ had the same dimensions as hemisphere-specific features, except all values were zero. (D) The template was generated using counterfactual brains from all participants, and then the Jacobi determinants between the left and right factual brains of each participant and the template were computed. We conducted paired t-tests on the two groups of Jacobian determinants representing structural changes induced by left- and right-brain-specific separately, followed by multiple comparison correction using 10,000 permutations.

## 2. Data and methods

### 2.1. Subjects

1,113 subjects with T1-weighted structural MRI from the public dataset S1200 Human Connectome Project Young Adult (HCP-YA) were used here. 47 participants were excluded because of the not good imaging quality. To avoid possible confounding effect of handedness, only right-handed HCP subjects whose Edinburgh Inventory Handedness score [24] exceeding + 40 were included (59 non-right-handed participants were excluded). As a result, 907 right-handed participants were included, with ages ranging from 22 to 36 (N_p_ = 907; 397 Males, 510 Females). All subjects provided written informed consent, and the research protocol was approved by the Institutional Review Board of Washington University. More specific scanning parameters please refer to [25].

### 2.2. Imaging acquisition and preprocessing

MRIs were acquired using a Siemens Connectome Skyra 3 T scanner with 32-channel head coil housed at Washington University in St. Louis [25]. For T1-weighted images, 256 slices per slab were acquired with a three-dimensional (3D) magnetization prepared rapid gradient echo sequence (MPRAGE). The detailed scanning parameters were as follows: TR = 2400 ms, TE = 2.14 ms, reversal time (TI) = 1000 ms, flip angle = 8^◦^, FOV = 224 × 224 mm^2^, resolution = 0.7 × 0.7 × 0.7 mm^3^, and total scan time = 7 min and 40s.

T1-weighted images were preprocessed using the HCP minimal-preprocessing pipelines [26]. In order to disentangles the hemispheric-specific feature, additional processing was executed to facilitate the implementation of a convolutional neural network. First, we split whole-brain T1 images into hemispheres images with the following steps: 1) align the whole-brain T1 images to the symmetric standard space by using the nonlinear transformation; 2) mask the whole-brain data with hemispheric brain mask; 3) transform the hemisphere brain imaging data back to native space; 4) mirror the right hemispheric images to the left ones for comparative analysis. Then, we made boundary cuts to the 3D images of the half brain, removing as much of the non-brain portion of the images as possible to minimize the learning of useless information during training. Finally, the hemispheric brain images were normalized to the range of 0 to 1 and resampled to a target resolution of 64 × 64 × 64.

### 2.3. In-Scanner Cognitive Performances

Referring to previous study [9], we selected two in-scanner cognitive performances to quantify the correlation between hemisphere features and cognitive performances. These tasks encompassed the HCP language processing and HCP social cognition performances [27], with language representing an anticipated left-brain dominant behavior and social cognition aligning with an expected right-brain dominant behavior. Language-related indicators included ‘average of accuracy from each condition in the language task’, ‘average of median correct reaction time from each condition in language task’, ‘accuracy percentage during story condition in language task’, ‘median reaction time for correct trials during story condition in language task’, ‘accuracy percentage during math condition in language task’ and ‘median reaction time for correct trials during math condition in language task’. Social-related indicators included ‘average of median reaction times from stimuli that the subject rated as “random” in social task’, ‘average of median reaction times from stimuli that received a “social” rating in social task’, ‘median reaction time for random stimuli that the subject rated as “random” in social task’ and ‘median reaction time for social stimuli that received a “social” rating in social task’ (N_langu_condi_ = 6, N_social_condi_ = 4).

### 2.4. Decoupled Representation Learning Model for Hemisphere Features

We designed a deep learning model named Fluffy for extracting hemisphere-specific and hemisphere-shared structure features based T1 weighted images. The model was built upon a contrastive Variational Autoencoder (cVAE) [28,29], an extension of the standard VAE [30] that allowed users to identify latent factors that were salient to their own analyses through the use of a background dataset. The Fluffy was composed by three probabilistic encoders and two probabilistic decoders (as shown in Figure 2). Encoders of Fluffy could effectively decouple 3D structural MRI data of the left and right hemispheres into mutually independent 16-dimensional hemispheric features (N_f_ = 16). These decoupled hemispheric features were then utilized by decoders to generate synthetic data that closely resembled the input data. Details regarding the objective function please find in the Supplementary materials. Notably, all encoders shared the same architecture, only the parameters of the neural network were different because of the different training data. The same was true for decoders.

**Fig. 2.**
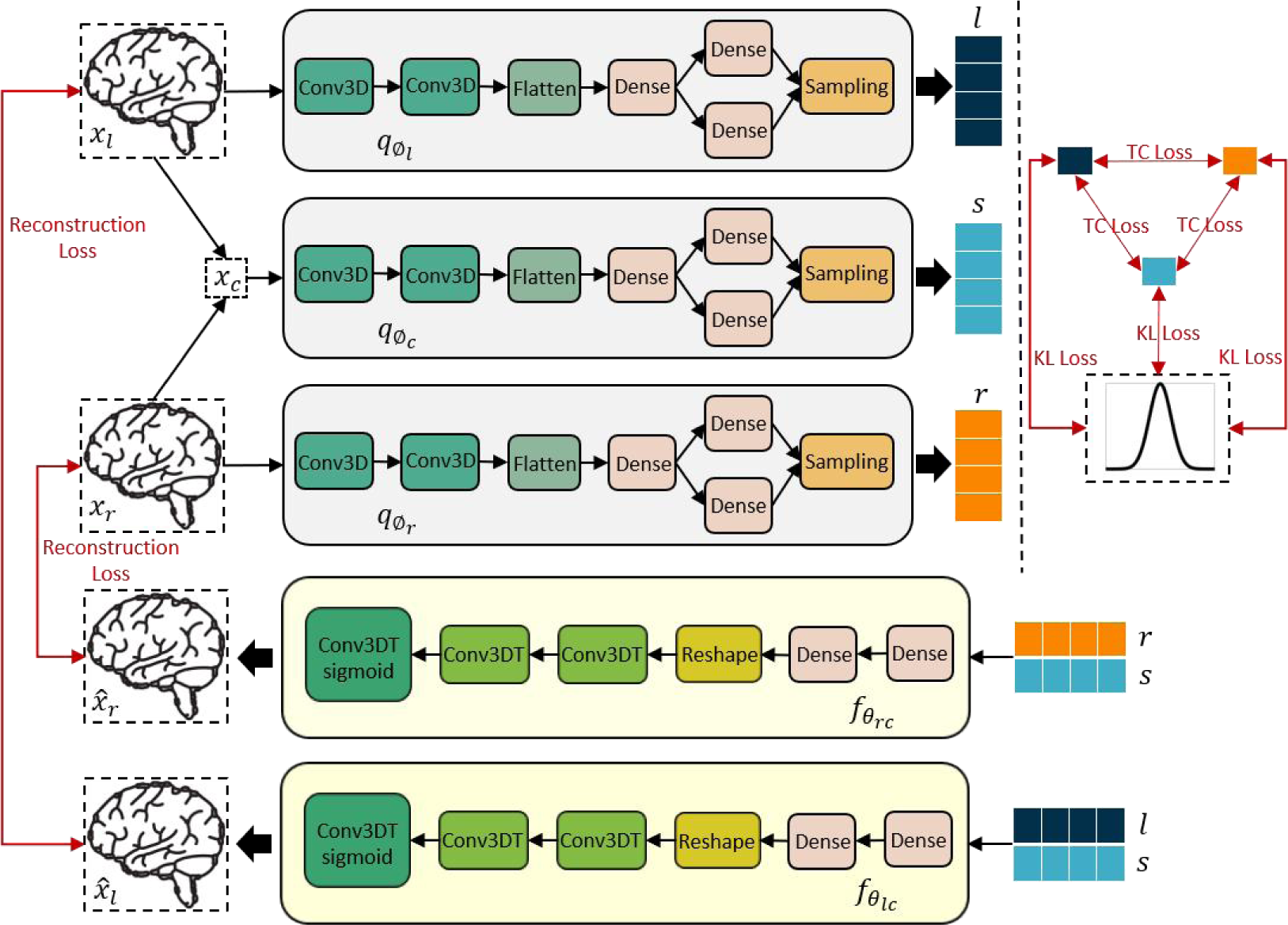
Architecture of Fluffy. ‘Conv3D’ is a three-dimensional convolutional layer; ‘Dense’ is a fully connected layer; ‘Conv3DT’ is a three-dimensional deconvolutional layer; ‘Conv3DT sigmoid’ is a three-dimensional deconvolutional layer with a sigmoid activation function. The red lines with bidirectional arrows represent the loss function used for optimizing the model. *x*, separated left and right brain structural data; 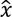, synthetic data generated by the model; *q*_θ_, encoders; *f*_θ_, decoders; *l*, left-specific features; *r*, right-specific features; *s*, shared features.

In the training phase, the model’s parameters were initialized randomly. The participants were divided randomly, with 80% assigned to the training set and the remaining participants to the validation set. The training process spanned 200 epochs. From the encoders, three distinct sets of hemispheric features were obtained: left-hemisphere-specific features, right-hemisphere-specific features, and shared features. These sets of features were intended for application in downstream tasks. Supplementary Fig.1. showed the loss curves during model training and the results of visualizing three sets of hemispheric features using the Uniform Manifold Approximation and Projection (UMAP) approach [31].

### 2.5. The correlation between hemisphere-specific features and the cognitive performances

In order to identify interpretable relationships between hemisphere-specific features and behavioral traits, Partial Least Squares Correlation (PLSC) [32,33] was employed to (Figure 1B). We hypothesized that if features captured by Fluffy contained hemisphere-specific information, left hemisphere-specific features would show a stronger association with language-related behaviors, while right hemisphere-specific features would demonstrate a stronger correlation with social behaviors.

As a multivariate technique, PLSC could capture major covariance patterns between two sets of data through singular value decomposition (SVD). This technique resulted in uncorrelated sets of latent variables (LV) which describe the linear combinations of the two datasets that were maximally covaried. Each LV represented a pattern of association between the two sets of data. In our study, the matrix for left/right hemisphere-specific features was N_p_ × N_f_, with N_p_ being the number of subjects and N_f_ denoting the dimensionality of hemisphere-specific features (N_p_ = 907, N_f_ = 16). Left-hemisphere-specific and right-hemisphere-specific features were concatenated into an N_p_ × (N_f_ + N_f_) matrix, serving as one of two input sets for PLSC. The other input set was one of two behavior matrices showing hemispheric dominance, with dimensions N_p_ × N_xxx_condi_, where N_xxx_condi_ represents the number of sub-behavioral traits within language/social behavior (N_langu_condi_ = 6, N_social_condi_ = 4).

To assess the statistical significance of output LVs, 10,000 permutation tests were performed, permuting rows of the hemispheric feature matrix (subjects) to obtain a null distribution of singular values. A non-parametric p-value for each LV was computed, applying a threshold of p < 0.05 for LV significance, signifying a 95% confidence that the singular value of the primary LV exceeds that of the permuted LV. To assess the contribution of each hemisphere feature to identified LVs, bootstrap resampling was employed. 1,000 sets of each hemisphere and behavior matrices were randomly sampled and rows replaced to create a distribution of singular vector weight. The bootstrap ratio (BSR) was calculated as the ratio of generated singular vector weight over the standard error of the weight from the bootstrap distribution. A threshold of p < 0.01 was applied for considering hemisphere features’ contribution significance (BSR < 2.58).

### 2.6. Contributions of hemisphere-specific features to structural changes

Tensor-based Morphometry (TBM) was employed to investigate the contributions of hemisphere-specific features to structural changes within the brain [34,35]. For each subject, we generated three synthetic brains: the first constructed from shared features and features with all zeros, termed the counterfactual brain; the second composed of left-hemisphere-specific and shared features, termed the synthetic left brain; and the third with right-hemisphere-specific and shared features, termed the synthetic right brain (Figure 1C). Here, ‘features with all zeros’ shared the same dimensions as hemisphere-specific features, except all values were zero. Subsequently, a reference template was created using counterfactual brains from all subjects (Figure 1D). Three sets of Jacobian maps were then obtained to characterize local volume differences between each subject’s three synthetic brains and the reference template.

To facilitate the identification of structural changes, we projected the gray matter portion of these Jacobian maps onto the Brainnetome Atlas (BNA), which consisted of 246 regions of interest (ROIs) [36]. The Jacobian maps from all subjects were then combined into matrixes of size 907 (number of subjects) × 123 (half of the number of BNA ROIs). Each value in these matrices represented the mean voxel-wise degree of brain structural volume change within a specific ROI of a certain subject. This step allowed us to conduct region-wise statistical tests between hemispheric Jacobian matrices within a common coordinate system. In each ROI, two-tailed paired t-tests were performed between region-wise hemisphere-specific and shared matrices to capture structural changes separately induced by the two hemisphere-specific features. Additionally, we employed permutation tests to assess the overall significance, correcting for multiple comparisons. A null distribution for the group differences in Jacobian matrices at each ROI was constructed using 10,000 random permutations. This permutation test approach allowed us to identify brain ROIs surpassing the significance threshold of p < 0.01. The final group-level t-maps, depicting significant structural changes in regional areas due to group differences, were generated by multiplying the masks derived from 10,000 permutation tests with the t-maps obtained from the preceding paired t-tests.

### 2.7. Relationship between significant changed ROIs and cognition

After obtaining the group-level t-maps illustrating structural changes linked to hemisphere-specific features, we conducted analyses using Neurosynth (https://neurosynth.org/) [37] to find the relationship between ROIs of brain gray matter undergoing structurally significant changes and cognitive terms. In this study, we utilized the “decoder” function to examine the relationship between left/right hemispheric t-maps and the meta-analytic map of each term in Neurosynth. The absolute values of the correlations obtained from “decoder” were taken, and the top 20 terms based on their absolute values were selected for presentation. Only the terms related to cognition in the results were retained, and singular and plural forms of the same word, such as “language” and “languages” were consolidated.

### 2.8. Gene expression associated with structural changes induced by hemisphere-specific features

To investigate the association between t-maps including all brain regions within one hemisphere and cortical gene expression levels, we performed Gene Enrichment Analysis (GEA). Specifically, gene expression data were obtained from Allen atlas which derived from 6 postmortem brains of donors without any neuropsychiatric or neuropathological diseases [38]. We projected gene data onto the BNA using the “abagen” toolkit (https://github.com/rmarkello/abagen/) and obtained 2 gene expression matrixes comprising 15,633 genes of 123 ROIs for left hemisphere and of 115 ROIs for right hemisphere (8 ROIs were excluded due to missing genetic data). Afterward, we computed the correlation between genetic data and the t-maps for each hemisphere, respectively. To strictly account for the spatial autocorrelation, we leveraged spin test to generate null distributions obtained by 1,000 spatial permutation tests, in which surrogate maps of t-maps were generated while maintaining the spatial autocorrelation properties of the original map. Only significantly expressed genes (p <0.05) were retained and utilized for GEA via the Metascape platform (https://metascape.org) [39].

### 2.9. Validation

In order to validate the correlation between hemisphere-specific features and the cognitive performances, we reran the PLSC analysis using a randomly selected two-thirds of the participants for language and social cognition.

## 3. Results

### 3.1. Fluffy successfully extracted hemisphere-specific features

The statistically significant LVs with p-values less than 0.05 were considered noteworthy. Each significant LV had hemisphere features with a BSR greater than 2.58 (p < 0.01), indicating a significant contribution to the LV. In terms of the relationship between hemisphere-specific features and language, a single significant LV explained 49.7% of the covariance (Figure 3A, p < 0.001). Notably, a left-specific feature demonstrated a significant contribution to the language-related LV after the robustness assessment (BSR = 3.462, p < 0.01). For the social aspect, the first LV explained 66.8% (p = 0.009) of the covariance. A right-specific feature significantly contributing to the social-related LV (BSR = 2.582, p < 0.01) was found (Figure 3A).

**Fig. 3.**
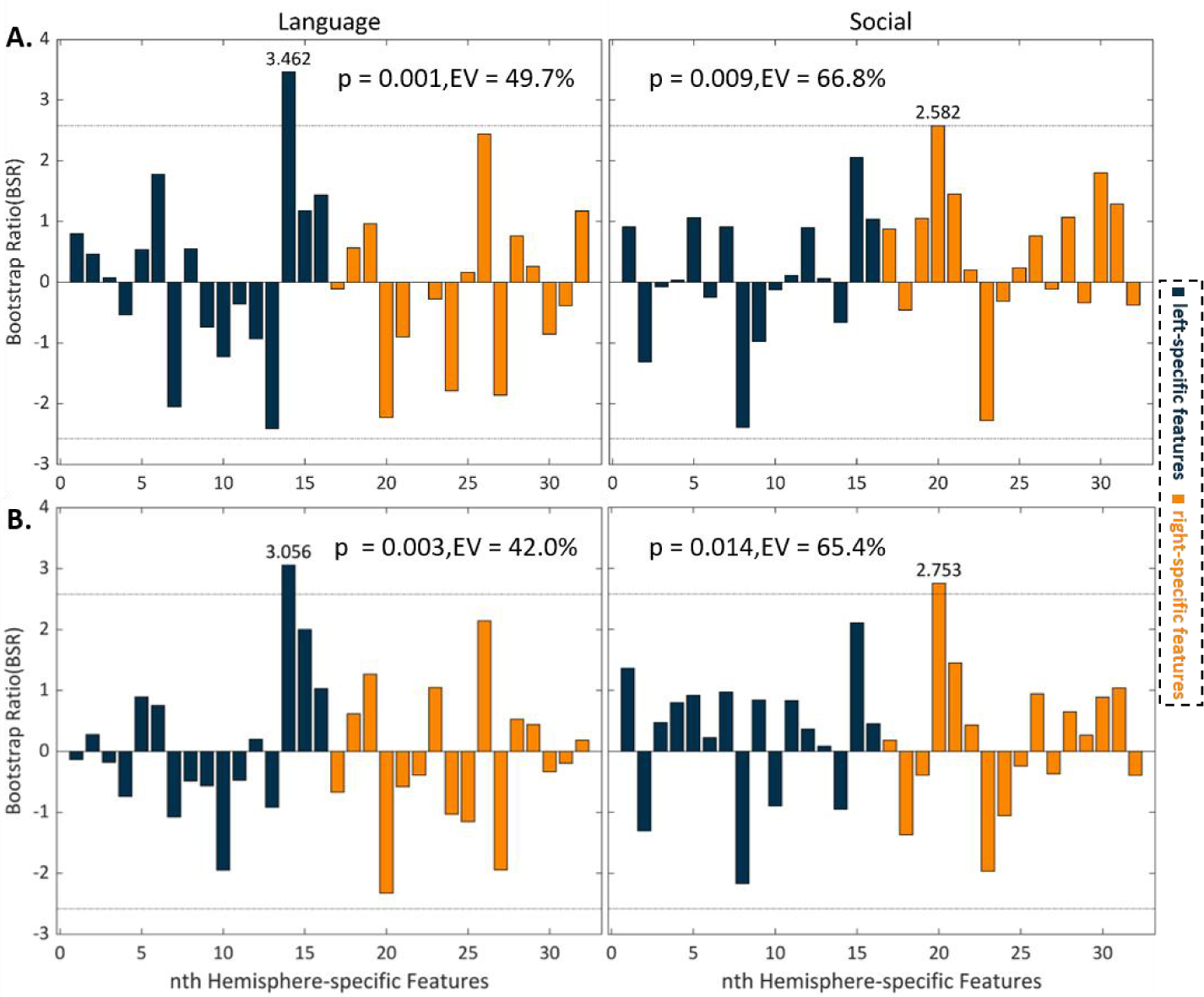
Results of PLSC. The figures displayed the significant latent variables. (A) The results of all participants. (B) Reproducibility verification. Redoing PLS results with a randomly selected 2/3 of the participants. The x-axis represented the order of hemisphere-specific features while the y-axis denoted the bootstrap ratio (BSR). The dashed line represented the significance threshold level. When BSR > 2.58 (p < 0.01), it indicated that the contribution of this feature to the current LV was significant. EV, explained variance.

The reproducibility validation, using a randomly selected 2/3 of the participants, confirmed the stability of our findings (Figure 3B). A single significant LV explaining 42.0% (p = 0.003) was found for language, with the same left-specific feature as the main results significantly contributing (BSR = 3.056, p < 0.01). In the social reproducibility results, the significant LV explaining 65.4% (p = 0.014) was identified, and the identical significant right-specific feature (BSR = 2.753, p < 0.01) was observed. All these results indicated that our decoupled representation learning model successfully extracted features containing hemisphere-specific information.

### 3.2. ROIs with significant structural changes were found

Figure 4A showed the significant gray matter ROIs of structural changes associated with hemisphere-specific features. ROIs with positive values indicated structural volume expansion of synthetic brains compared to the counterfactual brains, while negative values indicated volume compression. For left hemisphere, the main regions with significant expansion were lateral pre-frontal thalamus (lpfTha_l), caudal area in superior parietal lobule (cSPL_l) and middle postcentral gyrus (mPoG_l), while the regions with significant compression in volume were mainly located in inferior frontal sulcus (IFS_l), tongue and larynx region in precentral gyrus (tlPrG_l), rostral parahippocampal gyrus (rPhG_l), rostroventral inferior parietal lobule (rvIPL_l) and dorsal agranular insular gyrus (daINS_l).

**Fig. 4.**
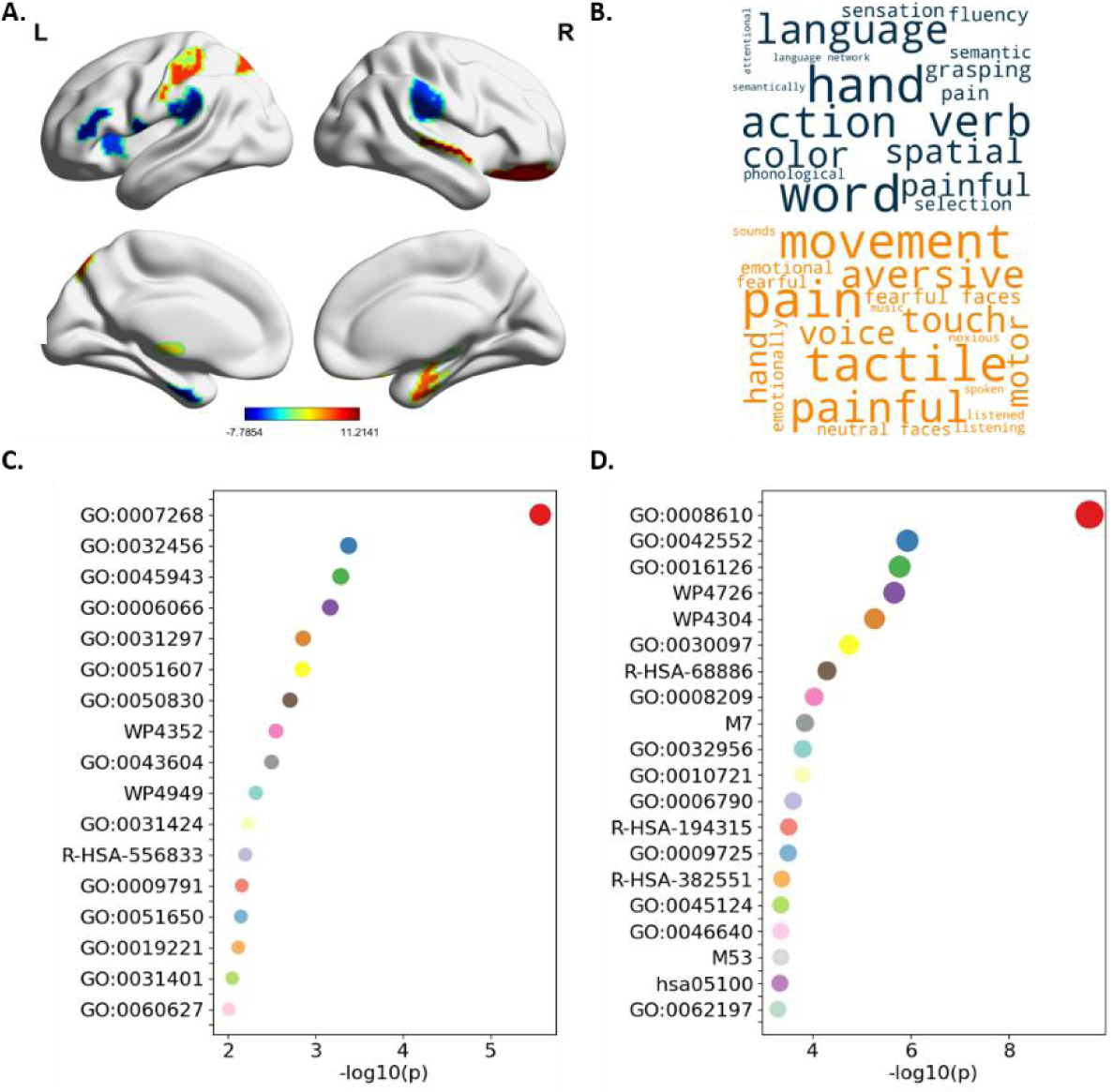
Results of TBM. (A) T maps depicting structural changes associated with hemisphere-brain-specific features. The first row was the lateral view, and the second row was the medial view. T map associated with left-hemisphere-specific features was in the left column, while t map associated with right-hemisphere-specific features was in the right column. The color intensity on the color bar indicated the magnitude of the values obtained from the t-test. (B) Word clouds of cognitive functions associated with t maps in (A). The font size of a given cognitive term corresponds to the correlation coefficient (r) generated by Neurosynth. (C) Results of GEA of left-brain-specific features. (D) Results of GEA of right-brain-specific features.

The map of right-hemisphere-specific features exhibited significantly expansion in rostral hippocampus (rHipp_r), lateral orbital gyrus (lOrg_r), rostral superior temporal gyrus (rSTG_r), rostral parahippocampal gyrus (rPhG_r), and dorsolateral putamen (dlPu_r) and significantly compression in rostroventral inferior parietal lobule (rvIPL_r) and sensory thalamus (sTha).

### 3.3. Word clouds showed typical cognitive terms with hemispheric dominance

As depicted in Figure 4B, the word clouds generated by Neurosynth revealed that the left-hemisphere t map exhibited correlation with various cognitive terms associated with language, such as “language”, “verb”, “semantic”, “word”, “semantically”, and “language network”. On the other hand, the right-hemisphere t map demonstrated correlation with terms like “pain”, “painful”, “emotion”, “emotionally”, “fearful”, “fearful faces”, and “neutral faces”, indicating a dominance of the right hemisphere in processing these aspects.

### 3.4. Hemisphere-specific features were associated with gene expression

After excluding the influence of spatial autocorrelation, 137 genes were significant correlation for left hemisphere and 254 genes were significant correlation for right hemisphere (p < 0.05). The GEA revealed that chemical synaptic transmission (GO Biological Processes, GO:0007268) showed the most correlation with t map of structural changes induced by left-brain-specific features. The post-embryonic development (GO Biological Processes, GO:0009791), representing the biological development process from the embryonic stage to maturity, also exhibited a significant correlation with left t-map (Figure 4C). For right hemisphere, lipid biosynthetic process (GO Biological Processes, GO:0008610) was the gene most associated with the specificity of the right brain. Myelination (GO Biological Processes, GO:0042552) was also one of the significant genes of right t-map (Figure 4D).

## 4. Discussion

By combining the decoupled representation learning and structural MRI, we systematically extracted hemisphere-specific structural features from the human brain. Our findings revealed that the hemisphere-specific multidimensional features, as extracted by the encoders of our model, exhibited correlations with lateralized behavioral performance (language and social). Then we utilized the model’s decoders to generate three sets of synthetic brains. Employing TBM, we identified several significant ROIs related to hemisphere-specific multidimensional features. In the left hemisphere, IFS_l and lpfTha_l, associated with language abilities, were pinpointed. Meanwhile, in the right hemisphere, rHipp_r, linked to visual and emotional processing, was identified. Neurosynth results highlighted that the left hemisphere-specific significant ROIs predominantly correlated with cognitive terms related to language, while the right hemisphere-specific ROIs correlated with emotional and facial recognition terms. Additionally, we found significant associations between t-maps including all brain regions and several genes, like gene of post-embryonic development and gene of myelination. Grounded in the hypothesis that the structure of the human brain comprises three factors and seamlessly integrating deep learning techniques, our study suggested that hemisphere-specific features could be disentangled through a data-driven approach to some extent. It is foreseeable that in the future, the fusion of artificial intelligence technology and brain asymmetry will become increasingly intertwined, paving the way for breakthroughs research.

For a significant duration in neuroscience, researchers faced constraints in conducting studies with small sample sizes, leading to heterogeneous conclusions on similar issues. However, over the last decade, the advent of large-scale datasets such as HCP, UK Biobank, Enhancing NeuroImaging Genetics through Meta-Analysis (ENIGMA), etc., has enabled the discovery of more universally applicable findings. Adequate data has also provided researchers with the conditions to leverage the flourishing development of deep learning technologies. In recent years, there has been a proliferation of articles utilizing deep learning for medical imaging research, spanning various domains within neuroscience [40,41,42,43]. However, the application of deep learning in the specific niche of studying hemispheric asymmetry in the human brain has been relatively scarce. Our exploratory research aims to fill this gap, and to the best of our knowledge, our work represents the first application of decoupled representation learning for feature extraction from structural MRI data into the study of brain asymmetry. Besides, we derived hemisphere-specific features by examining cortical gray matter, subcortical structures, and white matter tracts at the systematic level of hemispheres. This approach shifted studies on brain structural asymmetry from a regional to a holistic perspective, from a univariate to a multivariate approach. Furthermore, the research on the differences between the left and right brain hemispheres has been a relative concept. Our analytical approach allows us to explore the structural specificity of the left and right hemispheres without relying on relative perspectives.

### 4.1. Brain structural and functional asymmetry

The results of PLSC using hemisphere-specific features and language behavior were in line with our initial expectation that left hemisphere-specific features would show a stronger association with language-related behavior. Neurosynth results of left hemisphere encompassed numerous terms related to language functions, such as “language”, “verb”, “semantic”, and “word”. These terms covered various aspects of language behavior. The term “verb” introduced the concept of “verb argument structure processing”, referring to the process of handling the structure of verbs and their related arguments (subjects, objects, etc.) in language processing, particularly in understanding sentences and phrases. The term “semantic” represented the meaning of language units and their interpretation in specific contexts. Semantics not only focused on the literal meaning of words but also considered their implicit meanings and the influence of context. The term “word” denoted the basic units of language, including nouns, verbs, adjectives, etc.

In the study of structural brain asymmetry, Broca’s area was a key focus. Our left-hemisphere TBM results highlighted IFS_l within Broca’s area after multiple comparison correction. Additionally, lpfTha_l, located in the thalamus, was identified, and previous studies suggested the thalamus’s involvement in cognitive operations related to speech production [44]. Furthermore, tlPrG_l, a region in the precentral gyrus associated with the tongue and larynx, demonstrated significance through rigorous statistical tests. cSPL_l in the superior parietal lobule was suggested to play a crucial role in counting, calculation paradigms. This kind of mathematical ability was often considered a manifestation of language skills [45].

As for the right hemisphere, our PLSC analysis focused on its correlation with social behavior. This choice was motivated by the absence of cognitive behaviors related to visual spatial processing in HCP In-Scanner Task Performance. Also, Rajimehr’s work highlighted social tasks involving visual stimuli of HCP had the dominance of in the right hemisphere. In our Neurosynth results, the term “social” emerged, although it didn’t rank in the top 20. The top 20 Neurosynth results revealed numerous terms associated with “pain” and “face,” consistent with prior studies emphasizing results aligning with the right hemisphere [10,11]. Additionally, terms like “voice,” “listened,” “listening,” and “sound,” related to auditory functions, featured prominently associated with the right hemisphere. Despite both hemispheres possessing auditory abilities, the right hemisphere was generally considered more specialized in the processing of pitch and music [46].

A notable finding in the right-hemisphere TBM results was rHipp_r, a part of the hippocampus associated with emotion regulation and spatial navigation. Another interesting finding was lOrg_r, located in the orbital gyrus, which was linked to emotion. Additionally, rSTG_r in the superior temporal gyrus was associated with music and audition, while rvIPL_r in the inferior parietal lobule was related to pain. It is noteworthy that the Neurosynth results for the right hemisphere were consistent with the TBM findings. In the context of PLSC results, it is suggested that if there were tasks relevant to visual-spatial aspects in the In-Scanner Task Performance, our right hemisphere-specific features may exhibit significant correlations with them.

In our TBM results, we also identified several brain regions not directly associated with left or right brain dominance functions. Notable regions in the left brain included rPhG_l in the parahippocampal gyrus, related to memory; rvIPL_l in the inferior parietal lobule, and mPoG_l in the postcentral gyrus, associated with execution functions. In the right brain, significant regions included rPhG_r in the parahippocampal gyrus, as well as two subcortical nuclei, dlPu_r in the dorsolateral putamen of the basal ganglia, and sTha in the thalamus, both linked to execution. Our approach utilized deep learning technology to extract hemisphere-specific features and morphological methods to capture ROIs with significant changes related to hemisphere-specific features, leading to novel findings. However, the specific functions of these brain regions within the hemisphere require further exploration in subsequent research.

In our final work, we delved into the associations between t-maps representing group differences and gene data. Previous studies have proposed genes linked to left-right brain hemisphere asymmetry and lateralization [47,48,49]. We found gene of chemical synaptic transmission showed the most correlation with t map of structural changes induced by left-hemisphere-specific features and gene of lipid biosynthetic process was most associated with right-hemisphere-specific features. By contrasting our findings with existing researches, our study offered a new perspective on the study of brain lateralization and its correlation with genes.

### 4.2. Deep learning technologies

Our model, Fluffy, integrates the theoretical foundations of generative models with the principles of contrastive learning to extract hemisphere-specific features from structural MRI of healthy individuals. In the field of computer vision, several generative models are employed, with one of the most renowned models being the Variational Autoencoder (VAE), which serves as the foundational deep learning model in this article. VAE is a probabilistic model where the encoder maps input data into a regularized latent distribution, and the decoder generates new data from this latent distribution. Additionally, two widely applied generative models are the Generative Adversarial Network (GAN) [50] and Denoising Diffusion Probabilistic Models (DDPM) [51]. GAN comprises a generator and a discriminator. The generator produces synthetic data, while the discriminator distinguishes between real and synthetic data. Through adversarial training, the generator and discriminator continually optimize their parameters. When the discriminator can no longer differentiate between real and synthetic data, GAN achieves a balanced state where the generator can produce data resembling real data. DDPM’s training involves forward-noising and backward-denoising stages. In the forward stage, small Gaussian noise is added to the original data in each step, gradually transforming it into standard Gaussian noise. In the backward stage, useful information is progressively recovered from the noise. Compared to VAE, GAN produces higher-quality samples and does not require assumptions about the hidden distribution. However, GAN’s training instability can lead to mode collapse and oscillation issues. DDPM appears to be a promising model, but its training demands significant computational resources, posing challenges when applied to large-scale 3D datasets. In the future, exploring brain lateralization using GAN and DDPM presents a promising avenue for research.

Contrastive learning seeks to uncover the intrinsic structure and features of data by maximizing the similarity between positive samples and minimizing the similarity between negative samples. Positive samples represent those we anticipate being as similar as possible to the target sample, whereas negative samples are those we expect to be as dissimilar as possible. In Fluffy, our aim is for left-hemisphere-specific features, right-hemisphere-specific features, and shared features to exhibit maximum dissimilarity, treating them as negative samples to each other. Recent breakthroughs in contrastive learning, exemplified by MoCo [52] and SimCLR [53], have marked significant advancements. Exploring applications inspired by these model paradigms is also slated as one of our forthcoming research objectives.

### 4.3. Interpretability of the extracted features

Central to this study is the effectiveness of utilizing Fluffy to untangle the factors contributing to structural brain asymmetry. Encoders compress the structural data input into low-dimensional representations, known as hemisphere features. Decoders, in turn, nonlinearly reconstruct these low-dimensional representations back into the original high-dimensional space. Since these hemisphere features are discovered in a data-driven manner, attempting to interpret these features as specific physiological processes or cognition is challenging. Using these features for downstream tasks is a straightforward approach to uncover the relationship between discovered features and physiological processes. Jung-Hoon Kim, for instance, employed VAE to unravel the generative factors influencing resting state activity. Subsequently, these identified features were utilized to delineate individual variations and facilitate individual identification [22]. Aidas Aglinskas utilized a contrast VAE to extract ASD-specific and shared neuroanatomical features in patients with Autism Spectrum Disorder (ASD). He conducted a representational similarity analysis on both nonclinical and clinical characteristics of ASD participants [23]. In our study, applying PLSC did reveal associations between certain hidden features and language/social behaviors. Nevertheless, it is important to exercise caution when interpreting these results using post-hoc analysis. For more robust conclusions, future research endeavors should delve into studying the causal relationships between the latent features learned from brain imaging data and physiological processes or behaviors.

We do not anticipate a one-to-one correspondence between hemisphere-specific features and physiological processes, demographic indicators, or cognitive behavioral metrics. like “respiratory rhythms”, “age”, “health status”, “language “ since these factors may be interrelated and exhibit coupling. Creating a model that can achieve complete decoupling is indeed a challenging task. Instead, a one-to-many correspondence is more reasonable to expect. For example, in the outcomes of the left hemisphere PLSC analysis, we discovered a feature strongly correlated with language behavior. Nevertheless, it is possible that this particular feature still maintains some level of association with other factors that we have yet to investigate. Furthermore, the hemisphere-specific features extracted in our study may not necessarily exclusively represent brain asymmetry; they could potentially signify other relationships, such as unique features specific to either the left or right hemisphere without direct inter-hemispheric comparison. These aspects await further exploration in our future investigations.

### 4.4. Limitations

Due to our relatively weak model architecture and limited computing resources, the efficacy of our hemispheric multi-dimension features extraction analysis is somewhat constrained. Looking ahead, there is potential to incorporate more sophisticated models with much stronger decoupling capabilities, leveraging more powerful computing resources to train on higher resolution and larger datasets. Also, our focus has been on structural MRI T1 data from healthy individuals, yet the research has the potential to extend into the exploration of lateralization in individuals with neurodegenerative diseases, mental disorders, neurodevelopmental disorders, and more. Furthermore, the inclusion of medical imaging modalities such as computed tomography (CT), positron emission tomography (PET), diffusion tensor imaging (DTI), and functional Magnetic Resonance Imaging (fMRI), could enrich our investigations.

## References

1. Broca, P. Remarques sur le siège de la faculté du langage articulé, suivies d’une observation d’aphémie (perte de la parole). Bulletin et Memoires de la Societe anatomique de Paris, 6, 330–357 (1861).

2. Wernicke, C. Der aphasische Symptomencomplex: eine psychologische Studie auf anatomischer Basis. Cohn & Weigert (1874).

3. Sperry, R. W. Cerebral Organization and Behavior: The split brain behaves in many respects like two separate brains, providing new research possibilities. Science 133, 1749–1757 (1961).

4. Gazzaniga, M. S. Forty-five years of split-brain research and still going strong. Nat Rev Neurosci 6, 653–659 (2005).

5. Toga, A. W. & Thompson, P. M. Mapping brain asymmetry. Nat Rev Neurosci 4, 37–48 (2003).

6. Knecht, S. Handedness and hemispheric language dominance in healthy humans. Brain 123, 2512–2518 (2000).

7. Mazoyer, B. et al. Gaussian Mixture Modeling of Hemispheric Lateralization for Language in a Large Sample of Healthy Individuals Balanced for Handedness. PLoS ONE 9, e101165 (2014).

8. Vigneau, M. et al. Meta-analyzing left hemisphere language areas: Phonology, semantics, and sentence processing. NeuroImage 30, 1414–1432 (2006).

9. Rajimehr, R., Firoozi, A., Rafipoor, H., Abbasi, N. & Duncan, J. Complementary hemispheric lateralization of language and social processing in the human brain. Cell Reports 41, 111617 (2022).

10. Joseph, R. The right cerebral hemisphere: Emotion, music, visual-spatial skills, body-image, dreams, and awareness. J. Clin. Psychol. 44, 630–673 (1988).

11. Grand, R. L., Mondloch, C. J., Maurer, D. & Brent, H. P. Expert face processing requires visual input to the right hemisphere during infancy. Nat Neurosci 6, 1108–1112 (2003).

12. Hartwigsen, G., Bengio, Y. & Bzdok, D. How does hemispheric specialization contribute to human-defining cognition? Neuron 109, 2075–2090 (2021).

13. Steinmetz, H. Structure, Function and Cerebral Asymmetry: In Vivo Morphometry of the Planum Temporale. Neuroscience & Biobehavioral Reviews 20, 587–591 (1996).

14. Postema, M. C. et al. Altered structural brain asymmetry in autism spectrum disorder in a study of 54 datasets. Nat Commun 10, 4958 (2019).

15. Damme, K. S. F., Vargas, T., Calhoun, V., Turner, J. & Mittal, V. A. Global and Specific Cortical Volume Asymmetries in Individuals with Psychosis Risk Syndrome and Schizophrenia: A Mixed Cross-sectional and Longitudinal Perspective. Schizophrenia Bulletin 46, 713–721 (2020).

16. Kong, X.-Z. et al. Mapping Cortical and Subcortical Asymmetry in Obsessive-Compulsive Disorder: Findings from the ENIGMA Consortium. Biological Psychiatry 87, 1022–1034 (2020).

17. Roe, J. M. et al. Asymmetric thinning of the cerebral cortex across the adult lifespan is accelerated in Alzheimer’s disease. Nat Commun 12, 721 (2021).

18. Williams, C. M., Peyre, H., Toro, R. & Ramus, F. Comparing brain asymmetries independently of brain size. NeuroImage 254, 119118 (2022).

19. Saltoun, K. et al. Dissociable brain structural asymmetry patterns reveal unique phenome-wide profiles. Nat Hum Behav 7, 251–268 (2022).

20. Liu, H., Stufflebeam, S. M., Sepulcre, J., Hedden, T. & Buckner, R. L. Evidence from intrinsic activity that asymmetry of the human brain is controlled by multiple factors. Proc. Natl. Acad. Sci. U.S.A. 106, 20499–20503 (2009).

21. Kong, X.-Z. et al. Mapping cortical brain asymmetry in 17,141 healthy individuals worldwide via the ENIGMA Consortium. Proc. Natl. Acad. Sci. U.S.A. 115, (2018).

22. Kim, J.-H. et al. Representation learning of resting state fMRI with variational autoencoder. NeuroImage 241, 118423 (2021).

23. Aglinskas, A., Hartshorne, J. K. & Anzellotti, S. Contrastive machine learning reveals the structure of neuroanatomical variation within autism. Science 376, 1070–1074 (2022).

24. Oldfield, R. C. The assessment and analysis of handedness: The Edinburgh inventory. Neuropsychologia 9, 97–113 (1971).

25. Van Essen, D. C. et al. The Human Connectome Project: A data acquisition perspective. NeuroImage 62, 2222–2231 (2012).

26. Glasser, M. F. et al. The minimal preprocessing pipelines for the Human Connectome Project. NeuroImage 80, 105–124 (2013).

27. Barch, D. M. et al. Function in the human connectome: Task-fMRI and individual differences in behavior. NeuroImage 80, 169–189 (2013).

28. Severson, K. A., Ghosh, S. & Ng, K. Unsupervised Learning with Contrastive Latent Variable Models. AAAI 33, 4862–4869 (2019).

29. Abid, A. & Zou, J. Contrastive Variational Autoencoder Enhances Salient Features. (2019) doi:10.48550/ARXIV.1902.04601.

30. Kingma, D. P. & Welling, M. Auto-Encoding Variational Bayes. (2013) doi:10.48550/ARXIV.1312.6114.

31. McInnes, L., Healy, J. & Melville, J. UMAP: Uniform Manifold Approximation and Projection for Dimension Reduction. (2018) doi:10.48550/ARXIV.1802.03426.

32. Krishnan, A., Williams, L. J., McIntosh, A. R. & Abdi, H. Partial Least Squares (PLS) methods for neuroimaging: A tutorial and review. NeuroImage 56, 455–475 (2011).

33. Kalantar-Hormozi, H. et al. A cross-sectional and longitudinal study of human brain development: The integration of cortical thickness, surface area, gyrification index, and cortical curvature into a unified analytical framework. NeuroImage 268, 119885 (2023).

34. Lin, S.-J. et al. Neuroimaging signatures predicting motor improvement to focused ultrasound subthalamotomy in Parkinson’s disease. npj Parkinsons Dis. 8, 70 (2022).

35. Frackowiak, R. S. Human brain function. Elsevier (2004).

36. Fan, L. et al. The Human Brainnetome Atlas: A New Brain Atlas Based on Connectional Architecture. Cereb. Cortex 26, 3508–3526 (2016).

37. Yarkoni, T., Poldrack, R. A., Nichols, T. E., Van Essen, D. C. & Wager, T. D. Large-scale automated synthesis of human functional neuroimaging data. Nat Methods 8, 665–670 (2011).

38. Shen, E. H., Overly, C. C. & Jones, A. R. The Allen Human Brain Atlas. Trends in Neurosciences 35, 711–714 (2012).

39. Zhou, Y. et al. Metascape provides a biologist-oriented resource for the analysis of systems-level datasets. Nat Commun 10, 1523 (2019).

40. Yamins, D. L. K. & DiCarlo, J. J. Using goal-driven deep learning models to understand sensory cortex. Nat Neurosci 19, 356–365 (2016).

41. Zhuang, C. et al. Unsupervised neural network models of the ventral visual stream. Proc. Natl. Acad. Sci. U.S.A. 118, e2014196118 (2021).

42. Li, X. et al. BrainGNN: Interpretable Brain Graph Neural Network for fMRI Analysis. Medical Image Analysis 74, 102233 (2021).

43. Saxe, A., Nelli, S. & Summerfield, C. If deep learning is the answer, what is the question? Nat Rev Neurosci 22, 55–67 (2021).

44. Barbas, H., García-Cabezas, M. Á. & Zikopoulos, B. Frontal-thalamic circuits associated with language. Brain and Language 126, 49–61 (2013).

45. Gelman, R. & Butterworth, B. Number and language: how are they related? Trends in Cognitive Sciences 9, 6–10 (2005).

46. Gordon, H. Music and the right hemisphere. 3, 65-86). London: Academic Press (1983).

47. Thompson, P. M. et al. Genetic influences on brain structure. Nat Neurosci 4, 1253–1258 (2001).

48. Geschwind, D. H. & Miller, B. L. Molecular approaches to cerebral laterality: Development and neurodegeneration. Am. J. Med. Genet. 101, 370–381 (2001).

49. Geschwind, D. H., Miller, B. L., DeCarli, C. & Carmelli, D. Heritability of lobar brain volumes in twins supports genetic models of cerebral laterality and handedness. Proc. Natl. Acad. Sci. U.S.A. 99, 3176–3181 (2002).

50. Goodfellow, I., et al. Generative adversarial nets. Advances in neural information processing systems, 27 (2014).

51. Ho, J., Jain, A., & Abbeel, P. Denoising diffusion probabilistic models. Advances in neural information processing systems, 33, 6840–6851 (2020).

52. He, K., Fan, H., Wu, Y., Xie, S., & Girshick, R. Momentum contrast for unsupervised visual representation learning. In Proceedings of the IEEE/CVF conference on computer vision and pattern recognition 9729-9738 (2020).

53. Chen, T., Kornblith, S., Norouzi, M., & Hinton, G. A simple framework for contrastive learning of visual representations. In International conference on machine learning 1597–1607. PMLR (2020).

