## Supplementary Fig.1. for "Hemispheric multi-dimension features extraction analysis based on decoupled representation learning"

### Supplementary materials

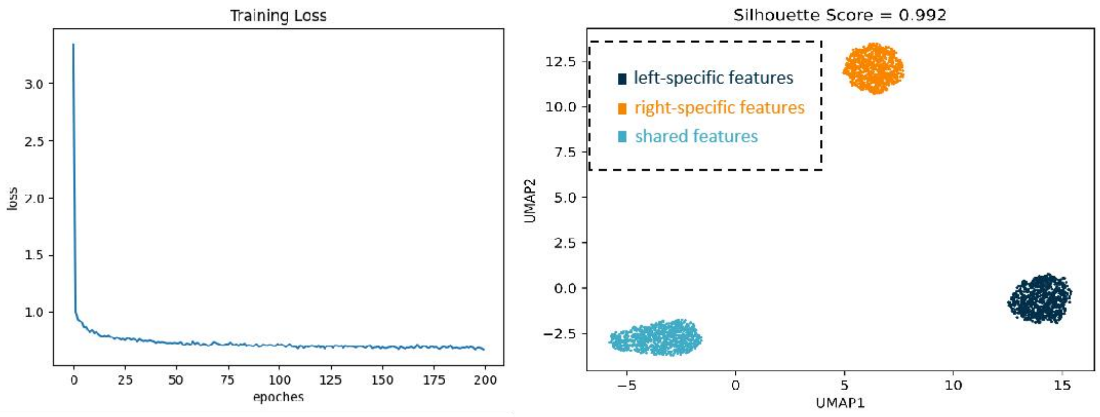


**Supplementary Fig.1.** Left: The loss curve during model training; Right: The results of visualizing three sets of hemispheric features using the Uniform Manifold Approximation and Projection (UMAP) approach. The silhouette score is indeed a metric used to evaluate the quality of clustering. It measures how similar an object is to its own cluster (cohesion) compared to other clusters (separation). The silhouette score ranges from -1 to 1, where a high value indicates that the object is well matched to its own cluster and poorly matched to neighboring clusters.
